# Stochastic gates for covariate selection in population pharmacokinetics modelling

**DOI:** 10.1101/2025.05.28.656586

**Authors:** Marija Kekic, Oleg Stepanov, Wenjuan Wang, Sam Richardson, Damilola Olabode, Carlos Traynor, Richard Dearden, Diansong Zhou, Weifeng Tang, Megan Gibbs, Andrzej Nowojewski

## Abstract

Covariate selection in population pharmacokinetics modelling is essential for understanding interindividual variability in drug response and optimizing dosing. Traditional stepwise covariate modelling is often time-consuming, compared to the new machine learning alternatives. This study investigates the use of Neural Networks with Stochastic Gates for automated covariate selection, aiming to efficiently identify relevant covariates while penalizing excessive covariate inclusion. On various synthetic datasets the approach demonstrated robustness in detecting important covariates, overcoming challenges such as high correlations, low covariate frequencies, high interindividual variability and complex covariate dependencies. In real clinical data from a monalizumab study, the method successfully identified covariates that matched those found by experts. However, for tixagevimab/cilgavimab, it identified a superset of covariates, indicating a potential need for further pruning. This machine learning-based method enhances the covariate pre-selection process in population pharmacokinetics model development, offering significant time savings and improving efficiency even under challenging scenarios.

**Graphical Abstract:** 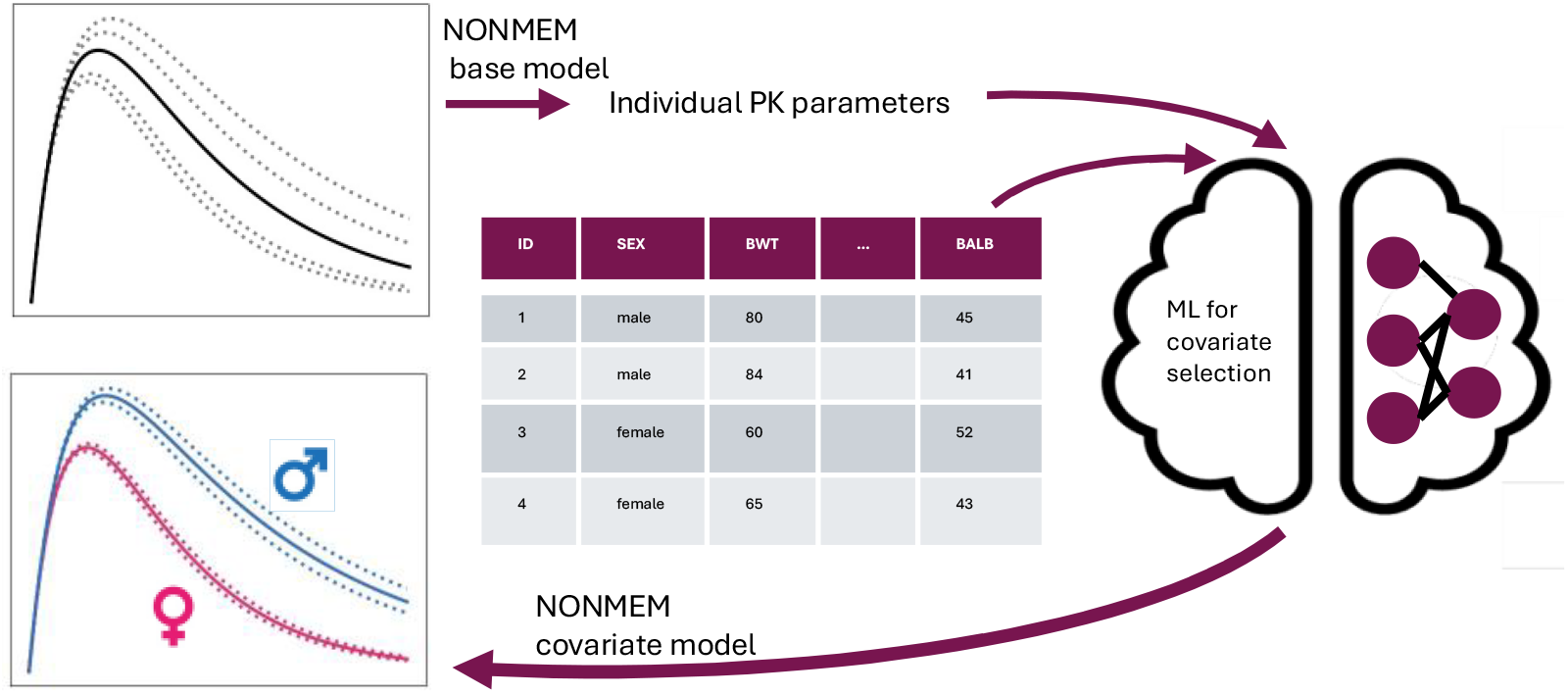

**Study Highlights:** *WHAT IS THE CURRENT KNOWLEDGE ON THE TOPIC?:* Covariate selection in population pharmacokinetics modelling is a crucial step in drug development and dosing determination. Traditionally, this process is conducted in a stepwise manner, which can be very time-consuming. Recently, fast machine learning (ML)-based methods for covariate selection have emerged in the literature, offering more efficient alternatives.

*WHAT QUESTION DID THIS STUDY ADDRESS?:* Can we use Neural Networks with Stochastic Gates, incorporating explicit penalization on the number of covariates, to select superior set of covariates compared to prior ML-based methods?

*WHAT DOES THIS STUDY ADD TO OUR KNOWLEDGE?:* Neural Networks with Stochastic Gates provide a reliable approach in accurately detecting covariates because they can effectively eliminate irrelevant covariates even in cases with high inter-covariates correlation.

*HOW MIGHT THIS CHANGE DRUG DISCOVERY, DEVELOPMENT, AND/OR THERAPEUTICS?:* This approach accelerates pharmacokinetic model development by significantly reducing the time required for covariate evaluation. Additionally, it allows for the screening of a wider set of covariates, potentially leading to fewer falsely missed covariates and better-quality models, a task that would be infeasible with traditional stepwise covariate modelling.

## Introduction

Developing a covariate model is an essential step in population pharmacokinetics (popPK) as it aims to understand the factors influencing interindividual variability^12^. The most extensively used covariate selection method in popPK is Stepwise Covariate Modelling^3,4^ (SCM) that relies on stepwise forward and backward inclusion of covariates and the differences in objective function (OFV) with and without included covariate. Optimised versions of SCM, such as SCM+^5^ or SAMBA^6^(Stochastic Approximation for Model Building Algorithm) include certain pruning of relations depending on the previous evaluations that allow for fewer estimation runs. Wald’s approximation^7^ is a method that relies on full covariate model covariance matrix estimation to approximate the differences in OFV between each reduced submodel and the full model, hence avoiding lengthy estimation steps occurring in SCM approach. LASSO^8,9^ (Least Absolute Shrinkage and Selection Operator) is an alternative approach proven superior to SCM for small datasets. Here, a penalty on the number of introduced covariates is added to the OFV, thereby reducing the number of introduced covariate-parameter relation.

Although all methods effectively identify relevant sets of covariates, finding the optimal set is never guaranteed, and none of the methods are well-suited for testing too many covariates. Additionally, covariate search is often conducted in the later stages of clinical trials, where populations typically include hundreds of individuals, making the iterative processes computationally infeasible due to the long estimation times. Therefore, pre-screening of covariates is often performed, relying on Empirical Bayes estimates of random effects (EBE). Common approaches include graphical correlations between EBE and covariates as well as the GAM^10^ (Generalized Additive Model) method. COSSAC^11^ (Conditional Sampling used for Stepwise Approach based on Correlation), a novel method that improves the speed of the SCM, also utilizes relationships between samples from individual random estimates and covariates prior to each SCM step.

The relationships between EBE and covariates have been further explored using machine learning (ML) to determine feature importance, aiming to identify covariates predictive of EBE variance among patients. In Ogami C et al. 2021^12^, an artificial neural network (NN) was used to model individual random effects based on covariates, with SHAP (SHapley Additive exPlanations)^13^ values employed to explain the model. In contrast, Janssen A et al. 2022^14^ employed an XGBoost^15^ model instead of a NN. Sibieude E et al. 2021^16^ conducted a systematic comparison of various tree-based and neural network models, using feature permutation techniques to rank covariates by importance. They also compared ML-based covariate selection with standard SCM and COSSAC, finding that novel ML approaches matched traditional techniques with substantial speed improvements. However, these methods do not provide explicit guidance on the number of covariates to use, and the authors suggest three possible strategies: top-M selection, order of importance (with a threshold for the sum of importances), and minimum degree of importance, each requiring a pre-specified choice of the number of covariates or importance threshold. Another alternative for pre-defined strategy is automatic recursive feature elimination based on SHAP values^17^.

In this study, we aimed to address the need for external feature removal and model retraining, by exploring an embedded feature selection method, known as Stochastic Gates^18^, for selecting relevant covariates based on EBE estimates. Unlike previous work, where covariate relevance is determined by SHAP or permutation tests, the Stochastic Gate method explicitly penalizes the number of input features facilitating model training and feature selection in a single step.

We evaluated the efficacy of the Stochastic Gates method through a series of synthetic data scenarios. These scenarios included datasets with correlated covariates of varying effect sizes, as well as challenging conditions of high intraindividual variability, imbalanced covariates, or highly nonlinear covariate dependencies. Additionally, we introduced a novel capability to assign preferences to certain covariates, as we demonstrated in a linearly dependent covariate scenario, allowing the method to prioritize certain covariates over others based on predefined criteria. Finally, we applied the method to two clinical datasets and compared with the pharmacometrician expert model.

## Methods

### Stochastic Gates for neural networks

Neural Networks^19^ (NN) are machine learning models that use layers of interconnected nodes to recognize patterns and model complex relationships in data. Yamada et al^18^ have recently proposed an extension of the LASSO penalization that is applicable to NNs, allowing for limiting the number of features used while maintaining full nonlinear modelling capacity.

The Stochastic Gates (SG) layer is a NN component placed immediately after the input layer that uses trainable parameters, *μ*_*d*_ “gates”, one per each input covariate, to probabilistically control the inclusion of each input feature. After the training, *μ*_*d*_ parameters are clipped to binary values (*z*_*d*_), which determine the inclusion of the covariate.

The training is done by minimising a loss function that composes of the mean squared error (MSE) loss, which is typically used in regression problems, and a penalty on the number of parameters. The penalty is scaled by a coefficient, *λ*, which governs the number of chosen features and should be carefully selected during hyperparameter optimization.

### Input data and training procedure

First, we fitted a base population PK model with intraindividual random effects and no covariates, to obtain Empirical Bayesian Estimates (EBE) using the nonlinear mixed effect framework in NONMEM^20^ (ICON Development Solutions, Ellicott City, MD)

In all the base models considered, we added the intraindividual random effects to the PK parameters using the standard log-normal distribution:

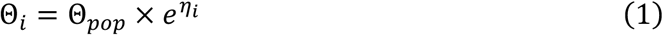

We then trained a NN to predict individual ETA (*η*_*i*_) parameters given patients’ covariates as the input (Figure 1).

**Figure 1:**
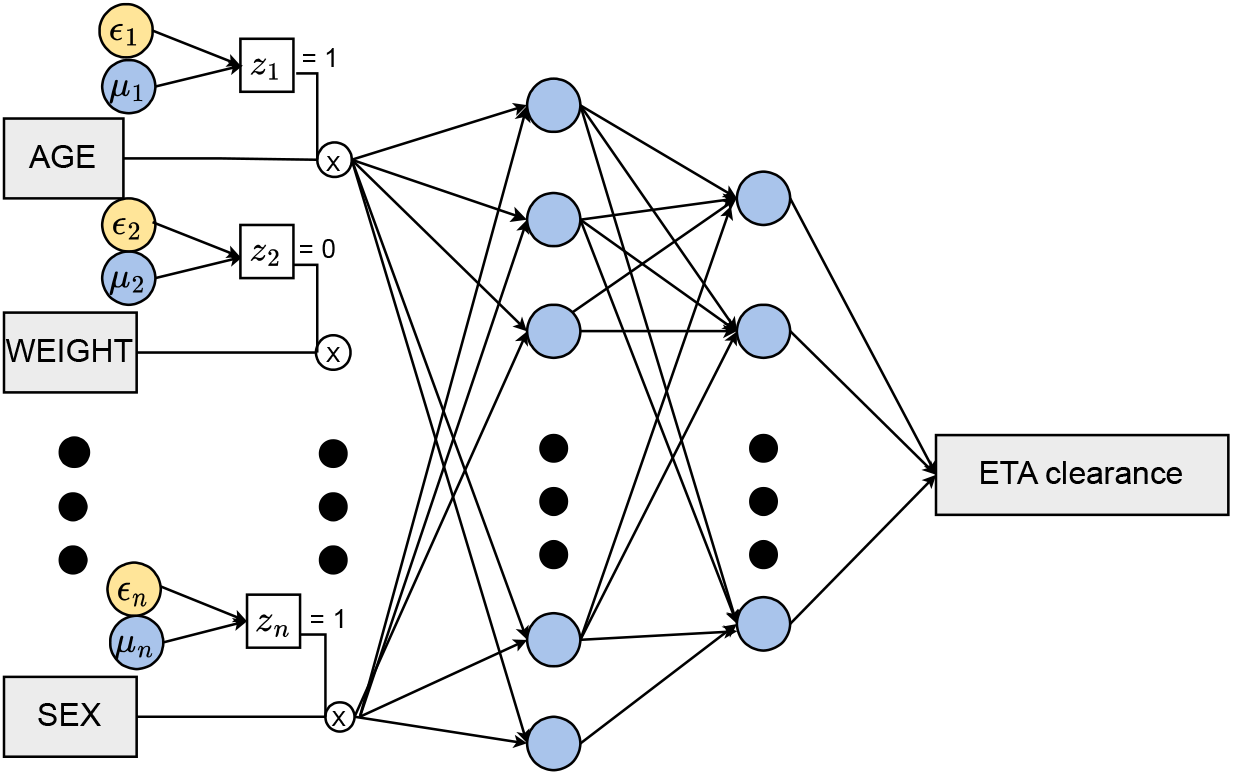
Schematic view of Neural Network with Stochastic Gate layer applied to covariate selection on the clearance parameter.

We divided the dataset into train and test, where 20% of the data is kept as the test data, with the training set used for hyperparameter tuning. For all our experiments, a two-layer NN with 20 hidden nodes in each layer and Adam optimizer with a constant learning rate of 0.001 were sufficient to fit the data well. To mitigate overfitting, we incorporated a dropout layer with a probability of 0.5 and a weight regularization term of 0.001.

To identify the optimal value of the crucial *λ* parameter, we performed a grid search across ranges 0.001 to 2 with 5-fold cross-validation (CV). We selected the optimal *λ* based on the validation loss observed during CV. Specifically, for each *λ*, we recorded the mean MSE validation loss, and the mean number of covariates utilized, choosing a *λ* that minimized MSE loss while using the fewest covariates. Additionally, we utilized a heatmap to illustrate the frequency of covariate selection across different CV folds for each *λ*. Features identified only in some folds may be due to noise rather than a true signal across the dataset. With hyperparameters fixed, we used the full training set to select the final set of covariates while monitoring performance on the test set.

Additional details on the method, input data preprocessing and hyperparameter tuning are provided in the Supplementary Material (S1).

### Warm start for preferred covariates

Sometimes certain covariates are preferred over others if their inclusion explains the data equally well, for example body weight might be preferred over body surface area. Such preferences can be implemented in the Stochastic Gates layer as initial values of the *μ*-gate parameter. Namely, instead of sampling from a random Bernoulli probability at the start of the training, *μ* can be set higher for preferred covariates and lower for the ones we disfavour. In our synthetic dataset experiments, setting priors in the range ±0.3 − 0.4 was successful in selecting preferred covariates.

### XGBoost and SHAP values

To compare with previous approaches, we trained an XGBoost model to predict individual *η* parameters from covariates and applied SHAP values to obtain feature importances. The SHAP method is a technique that evaluates feature sensitivity and derives an additive decomposition from the underlying data.

We tuned the hyperparameters of XGBoost on the validation set, provided final SHAP plots for the test set, and detailed the model training in the S1.

## Dataset

We first demonstrated the method by using a synthetic data and later applied it to real data from two studies: monalizumab and tixagevimab/cilgavimab.

### Synthetic data generation

For the synthetic data generation, we fixed the structural model to a two-compartment model with first order absorption and elimination in the single dose regimen. We constructed covariate models using standard formula for individual (*i*) with continuous (*CON*_*i,j*_) and categorical (*CAT*_*i,k*_) covariates:

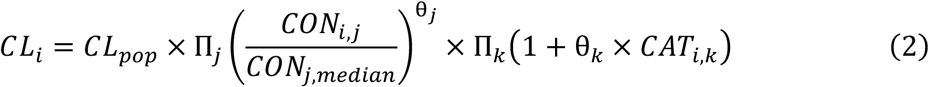

We sampled 1,000 patients from multidimensional normal distributions with 12 continuous and 4 categorical covariates. Correlations ranged (in absolute value) from 0 to 0.95, with the highest between BWT and BSA. Categorical covariates were binary with balanced frequencies (50%). We provided a complete list of covariates and their correlation plot in Figure S1.

To demonstrate the method, we simulated the following scenarios.

*Reference* scenario *with highly correlated covariates* included continuous covariates BWT and AGE with effect sizes 0.3 and 0.1 respectively, as well as two categorical covariates, SMK and COPD, with effect sizes 0.3 and 0.1. Effect size is defined as a ratio of the clearance parameter between the two stratified group (in case of categorical covariates), or a maximum between ratio of 5^th^ or 95^th^ percentile and the median value (in case of continuous covariates). We set intra individual variability (IIV) to 20% and chose the additive error model of ∼1% of the maximum PK concentration.

In the *Linearly dependent covariates* scenario, we demonstrated how warm-start affects the training by adding additional ‘cohort’ features, as a linear combination of two relevant covariates, SMK and COPD. Specifically, patients having negative SMK and COPD status belonged to the first cohort, patients with either positive SMK or positive COPD to the second and patients with both SMK and COPD positive to the third cohort. With this deliberately constructed relationship, the data can be described equally well by including any two of the three covariates mentioned, hence, without a feature preference method, the choice among these covariates would be random. This scenario simulated the effect of non-physiological covariates that may be selected in an automatic covariate search due to their correlation with biologically meaningful covariates, but an expert would deprioritize them in favour of those with biological significance.

Identifying covariates with low frequencies can be challenging, especially for those with small effect sizes. We investigated this in the *Low-frequency categorical covariates* scenario, where we simulated the dataset using the same base and covariate model as in the reference scenario, but reduced the frequency of both categorical covariates to 5% for the positive class.

In the *High Intra Individual Variability* scenario we increased the IIV from reference value to 120%. High IIV can act as noise obscuring relevant covariates in EBE-based covariate selection.

Finally, to illustrate the advantages of an ML approach compared to linear models, we simulated a *Highly nonlinear* scenario using a textbook nonlinear exclusive OR (*XOR*) function, described in S1. Univariate covariate screening would be unable to identify such covariate effects; hence such relationship would be missed in the standard EBE-based selection.

### Clinical datasets

*Monlizumab* is a first in class humanized IgG4 monoclonal antibody immune checkpoint inhibitor targeting NKG2A, administrated intravenously. Data from two clinical studies in patients with advanced solid tumors were pooled for the analysis, resulting in a dataset of PK samples from 507 subjects.

The structural model^21^ was identified as a two-compartment model with first-order elimination. PK parameters include clearance (CL), volume of distribution (Vc), peripheral volume (Vp), peripheral clearance (Q). In the base model, *η* values for CL and Vc exhibited relatively high correlation as presented in Figure S2a. Covariates explored included 9 continuous and 5 categorical covariates, as summarized in the original source.

Final expert model^21^ included baseline albumin levels and body weight as significant predictors of CL; body weight, sex, and smoking status significantly influenced Vc. However, none of the covariates were clinically significant.

*Tixagevimab/cilgavimab* is a combination of SARS-CoV-2-neutralizing antibodies developed for pre-exposure prophylaxis and treatment of COVID-19. PK data was pooled from 8 clinical trials with a total of 4,940 participants. The base model^22^ featured a two-compartmental distribution with first-order absorption and elimination, incorporating standard allometric exponents to account for the effect of body weight on clearance and volume. Concentrations were calculated as a sum of tixagevimab and cilgavimab concentrations.

PK parameters included clearance, volume of distribution, absorption rate (ka), peripheral volume, peripheral clearance and bioavailability, with IIV included on all but the bioavailability parameter.

Covariate analysis included 5 continuous and 14 categorical variables (see original publication for summary). Relatively high correlation between η values for CL, Vc and ka is demonstrated in Figure S2b.

Covariate model^22^ found by the expert included sex, advanced age (categorical), high body mass index (categorical), and diabetes affecting ka; diabetes influencing CL; Black race impacting Vc; and the intramuscular injection site affecting bioavailability. No covariates necessitated dose adjustment.

## Results

### Synthetic data

As discussed in the Methods section, to identify a suitable value for the gate penalization term λ, across wide λ range, we trained five CV models. We then evaluated the MSE loss as a function of λ and the number of active gates.

For the *Reference* scenario *with highly correlated covariates*, we found that the MSE loss reached a minimum when *λ* is within the range of 0.04 to 0.2, with exactly four covariates selected (Figure 2a). The reduction in the validation loss at this optimal number of input covariates, compared to when all covariates are used, is attributed to the network’s ability to avoid overfitting by not selecting excessive inputs that capture noise. The frequencies across different *λ* values (Figure 2b) indicated that the four discovered features are consistent across splits. Training and testing loss closely aligned with each other (Figure 2c), demonstrating that no overfitting occurred. In Figure 2d, we presented the convergence of the gate parameters (*μ*), where all the covariates whose *μ* values never rose are group together as the ‘Rest’. Covariates with *μ* values above the threshold line after convergence were considered relevant. Interestingly, at the onset of training, the *μ* parameter associated with BSA increased, indicating that BSA was initially deemed more relevant compared to other features due to its strong correlation with BWT. However, as training progressed, the importance of BSA declined, demonstrating the method’s effectiveness in handling scenarios involving highly correlated features. This decline occurred because BWT and BSA started competing against each other, leading to the selection of only one covariate.

**Figure 2:**
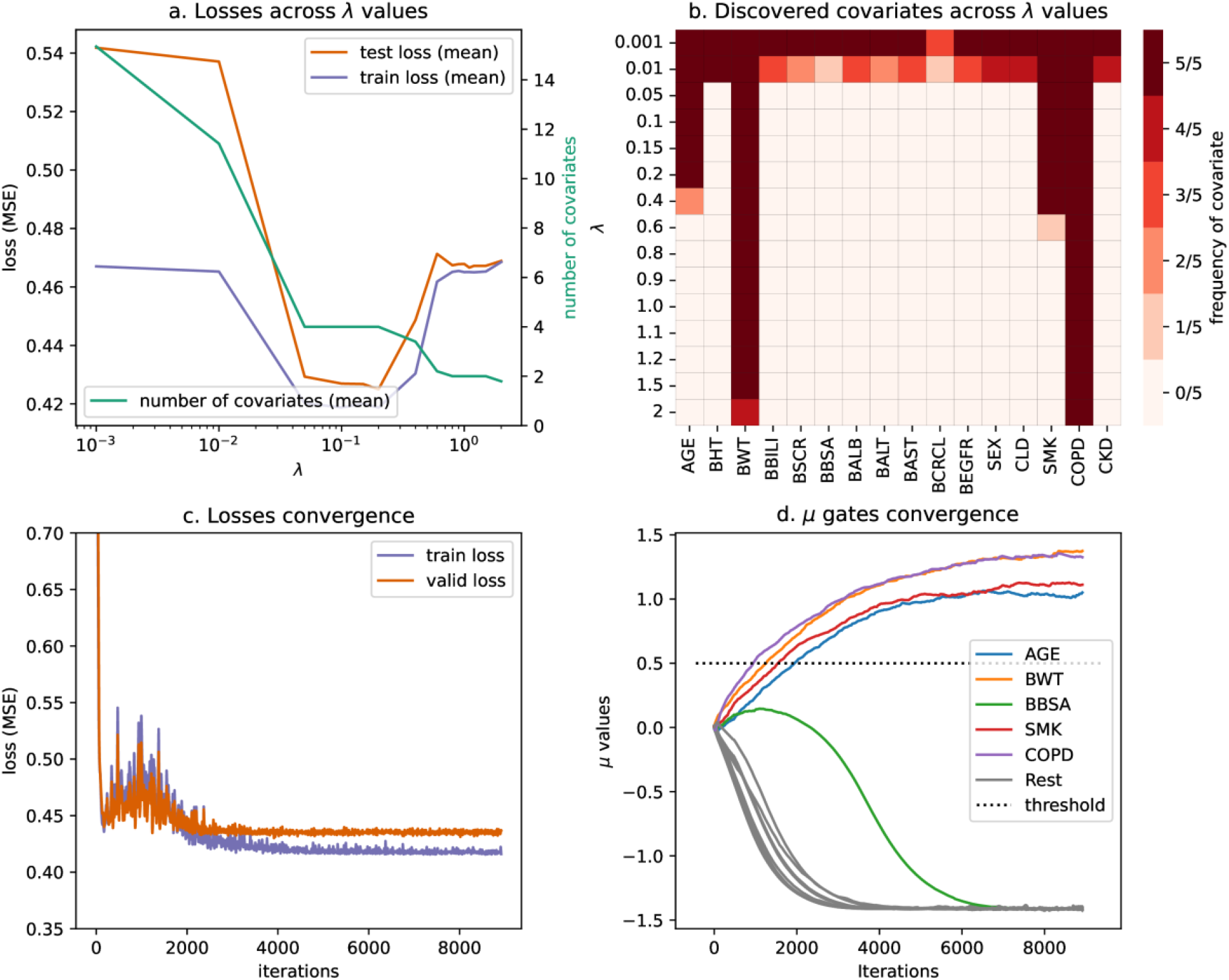
Stochastic Gates output for the Reference scenario. a) Validation and Training loss across different lambdas. b) Covariates frequencies across lambda values. c) Training and test loss convergence. d) *μ* gates convergence.

For comparison, we also conducted XGBoost modelling, utilizing SHAP values to rank features by their importance (**Error! Reference source not found**.). Although the top identified covariates were indeed the simulated ones, it is evident that XGBoost was unable to fully eliminate irrelevant covariates, resulting in ambiguity regarding the appropriate threshold for covariate inclusion.

**Figure 3:**
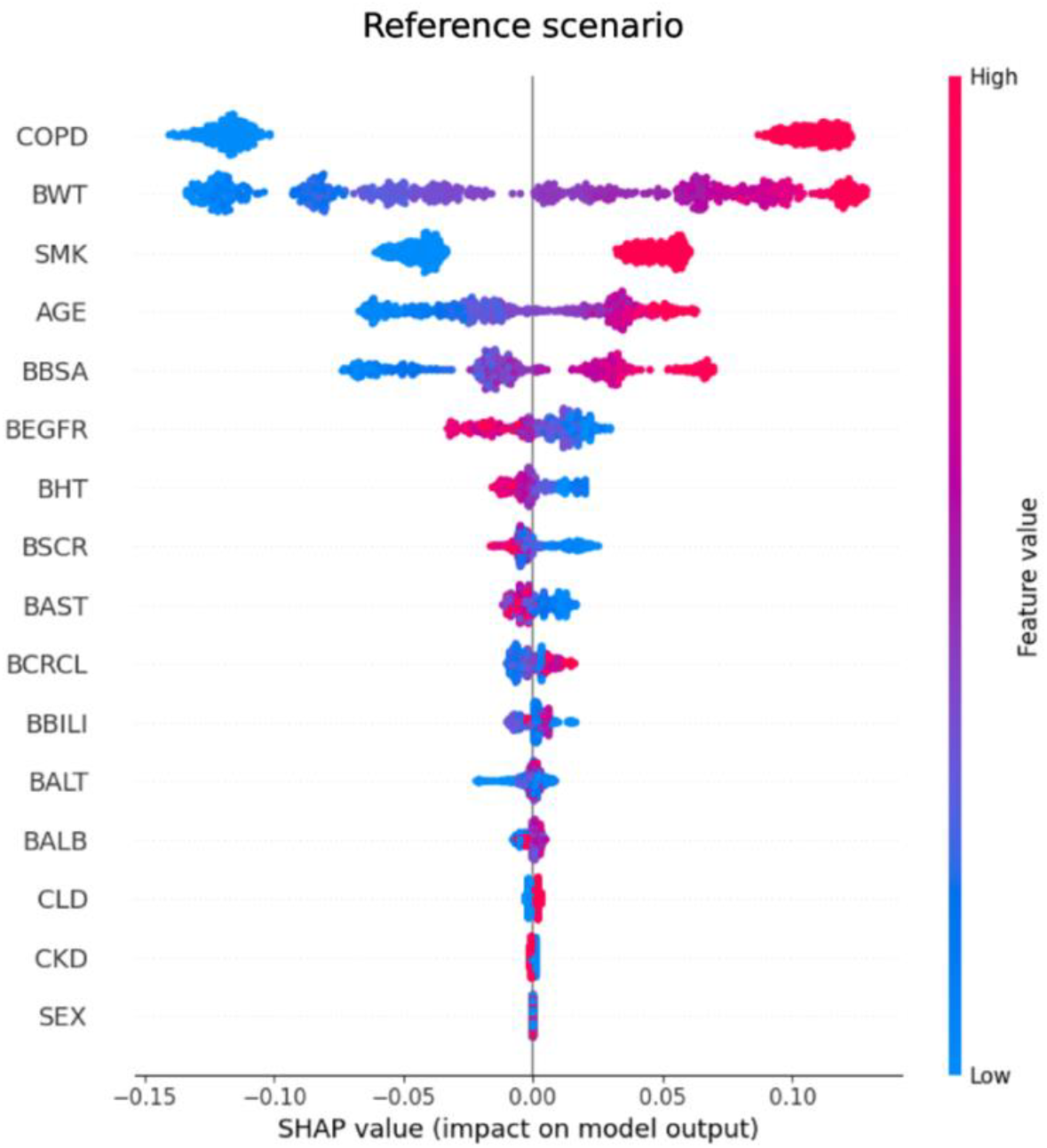
SHAP values of XGBoost model for the Reference scenario.

In the *Linearly dependent covariates* scenario, we demonstrated that the warm-start initialization can prioritize biologically relevant covariates (SMK and COPD) over the synthetic ‘cohort’ (linear combination of the two) covariate (see the Synthetic data generation section). Initially, we evaluated the *μ* gates without employing a warm start, resulting in the selection of the cohort while SMK was not chosen (**Error! Reference source not found**.a). Subsequently, we introduced a warm start by setting the *μ* gates to -0.3 for disfavoured covariates and 0.3 for favoured ones. Specifically, we gave preference to BSCR (not relevant) and disfavoured AGE (relevant) and the cohort (not relevant). In these settings, the algorithm successfully identified the four simulated covariates (**Error! Reference source not found**.b).

**Figure 4:**
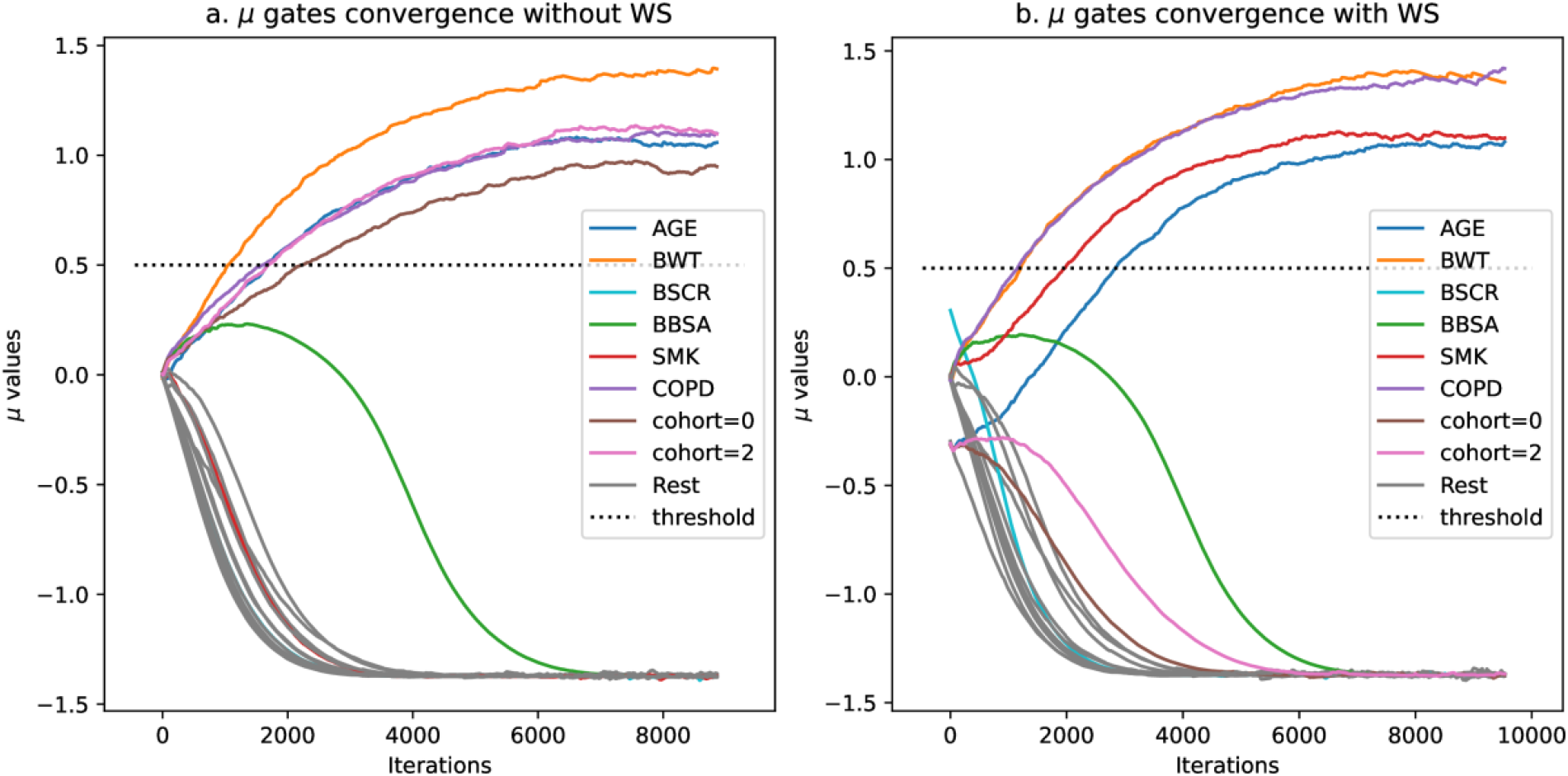
Convergence of the *μ* gates without (a) and with (b) warm start in the Linearly dependent covariates scenario.

We presented the resulting plots for the rest of the scenarios in Figure S3.

In the *Low-frequency categorical covariates* scenario only the medium effect covariate was discovered (COPD, 30).

In the *High interindividual variability* scenario, all four simulated covariates were successfully identified. However, the range of *λ* values over which the algorithm successfully detected these covariates was narrower. Additionally, the frequencies of the discovered covariates are less consistent compared to the easier, Reference scenario.

Finally, we presented the results for the *Highly nonlinear XOR* scenario, where the method successfully recovered the two non-linear covariates. Although the range of *λ* values for recovering these covariates was relatively narrow, the minimum validation loss was evident.

We provided the summary of the synthetic datasets and SG result in Table 1.

**Table 1:** Summary of synthetic dataset results.

| Dataset name | Description | Covariate Model | Discovered covariates |
| --- | --- | --- | --- |
| **Reference scenario with highly correlated covariates** | 'Easy' scenario with low IIV and balanced (but correlated) covariates | AGE, SMK (low effect)<br>BWT, COPD (medium effect) | AGE, SMK, BWT, COPD |
| **Linearly dependent covariates scenario** | 'Cohort' variable introduced to the Reference data, linearly dependent | AGE, SMK (low effect)<br>BWT, COPD (medium effect) | Without Warm Start:<br>AGE, BWT, COPD, cohort |
|  | on SMK and COPD |  | With Warm Start:<br>AGE, SMK, BWT, COPD |
| **Low-frequency categorical covariates** scenario | Imbalanced categorical covariates | AGE, SMK (low effect)<br>BWT, COPD (medium effect) | AGE, BWT, COPD |
| **High interindividual variability** scenario | Added 120% CV on CL | AGE, SMK (low effect)<br>BWT, COPD (medium effect) | AGE, SMK, BWT, COPD |
| **Highly nonlinear XOR** scenario | XOR covariate dependency | SMK, COPD | SMK, COPD |

### Clinical datasets

As with synthetic dataset, we repeated the procedure of *λ* search, recording validation loss and covariate frequencies.

For the *monalizumab* data, in **Error! Reference source not found**. we presented the training outcome of the two tested PK parameters (CL and Vc). Peripheral compartment parameters (Vp and Q) exhibited high shrinkage, hence, were not appropriate for EBE-based testing.

For the CL parameter, the validation loss reached a minimum with the inclusion of only two covariates: body weight and baseline albumin measurement. For the Vc, the minimum corresponded to a *λ* value around 0.2, which includes sex, smoke status, body weight, and albumin. The covariates identified by the model matched those selected by the expert, as summarized in Table 2. The precise covariate effects of the final expert model are detailed in the corresponding publication^21^.

**Figure 5:**
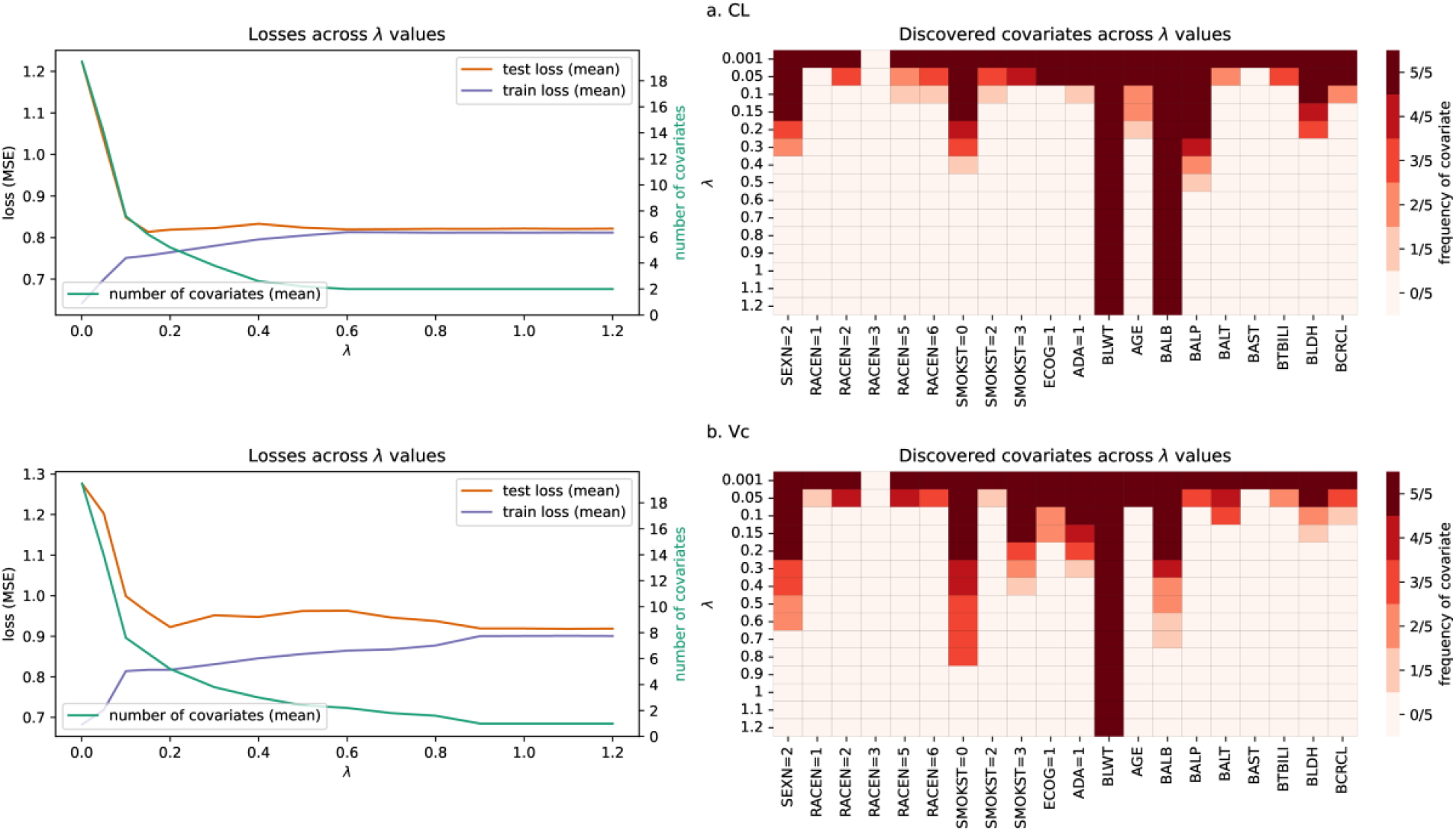
Monalizumab covariates for CL (a) and Vc (b). Left panels show validation/train loss and number of relevant features as a function of lambda parameter; right panels show frequency of chosen features across different lambda values

**Table 2:** Comparison of the expert model covariates and the covariates suggested by the SG method for the monalizumab dataset.

| PK parameter name | Expert model | Suggested covariates |
| --- | --- | --- |
| CL | BWT, BALB | BWT, BALB |
| Vc | BWT, BALB, SMK, SEX | BWT, BALB, SMK, SEX |

For the *tixagevimab / cilgavimab* data, we trained models for each of the three PK parameters: CL, Vc and ka (**Error! Reference source not found**.). Again, high shrinkage in the peripheral compartment parameters (Vp and Q) rendered them inappropriate for EBE-based testing.

For the ka, the model successfully identified all four covariates deemed relevant by the expert and additionally recognized the site of injection as relevant. In the expert model, the site of injection was not a covariate for the ka parameter but was a relevant covariate for bioavailability. For the Vc, the covariates found are sex, Black race, injection site, categorical age. In contrast, the expert model included only Black race as a parameter, suggesting that backward covariate elimination should be conducted before finalizing the model. For the CL, minimum validation loss corresponded to *λ* ≈ 0.1 and the set of approximately 8 covariates. However, by increasing the *λ* parameter to 0.3, the MSE loss increased from 0.89 to 0.91, while the number of covariates halved. In such a case, a modeler can choose higher *λ* to reduce the set of covariates, as the effect of dropped covariates should not be significant.

Main difference of proposed covariates with the expert model one is the repetition of certain covariates across all PK parameters, such as sex, that would require further pruning to achieve a minimal set of covariates. A possible cause for sex being detected on all PK parameters is the high correlation between random effects (around 40%, details in Figure S2b) and its large effect size (approximately 70% on the ka). For a detailed account of the covariate effects in the final model, please refer to the relevant publication^22^. We presented the full set of covariates identified by the expert and our model in Table 3.

**Figure 6.**
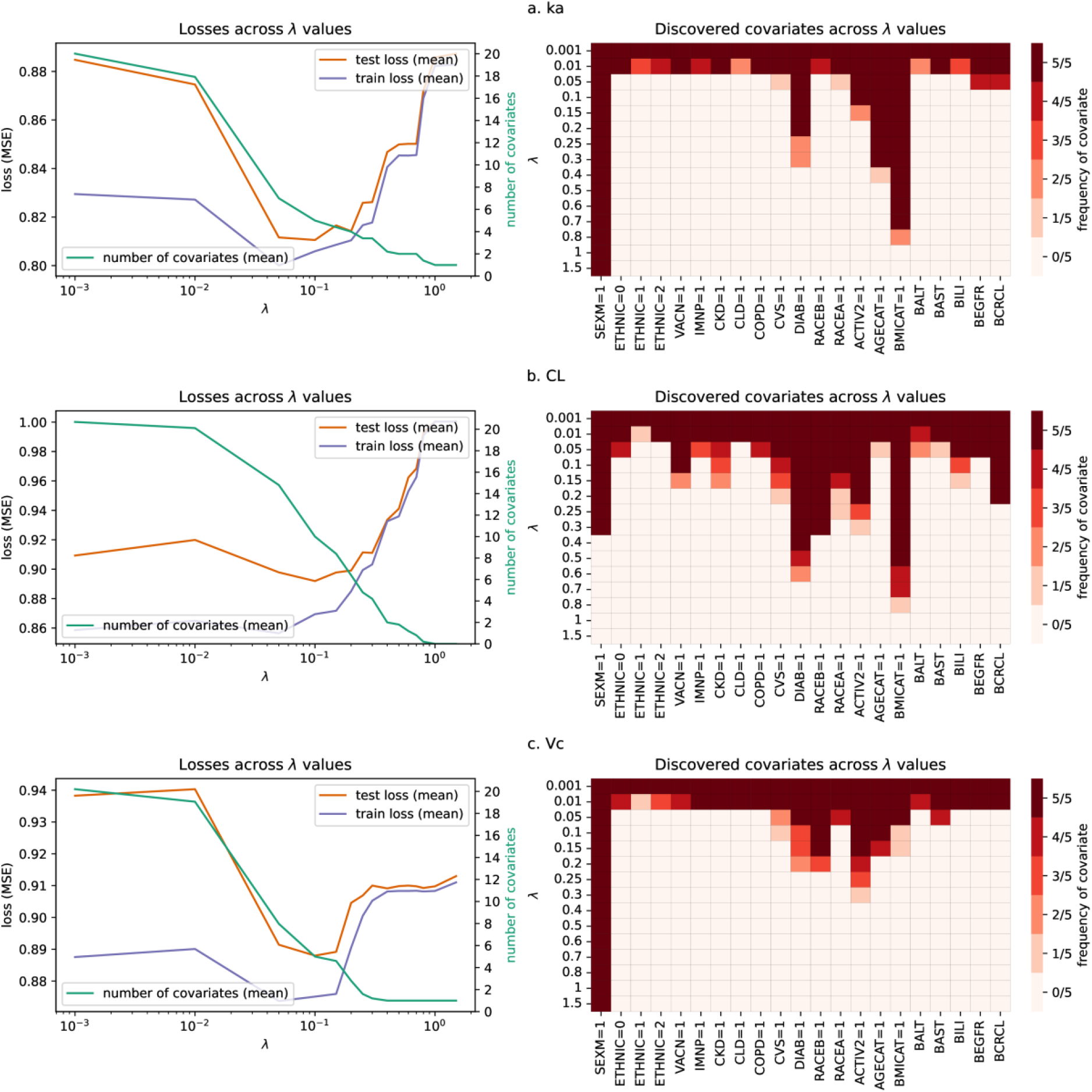
Tixagevimab / cilgavimab covariates for ka (a), CL (b) and Vc (c). Left panels show validation/train loss and number of relevant features as a function of lambda parameter; right panels show frequency of chosen features across different lambda values.

**Table 3:** Comparison of expert model covariates and covariates suggested by SG method for tixagevimab / cilgavimab dataset.

| PK parameter name | Expert model | Suggested covariates |
| --- | --- | --- |
| CL | Diabetes | Diabetes, Sex, Black race, BMI |
| ka | Categorical age, categorical BMI, sex, diabetes | Categorical age, categorical BMI, sex, diabetes, injection site |
| Vc | Black race | Sex, Black race, injection site, Categorical Age |
| bioavailability | Injection site | None (no IIV included) |

We give the final train/test losses and gate convergence for both clinical datasets in Figure S4.

### Comparison to XGBoost modelling

For all datasets, we used XGBoost and SHAP values to compare the identified covariates. We provided plots for all synthetic datasets, except for the reference scenario in Figure S5. The model often struggled to remove BSA from the top covariates, as well as having an unclear cut on the number of relevant covariates, since many had similar SHAP values. In the *Low-frequency* covariates scenario, even medium-effect ones were not among the top selected. Finally, the model was unable to prioritize biologically meaningful covariates in the *Linearly dependent* scenario. For both clinical datasets (Figure S6), covariates were consistently selected as in SG and by the expert. Performance metrics on training and test data were similar between SG and XGBoost, as shown in Table S1.

## Discussion

Previous studies have demonstrated some advantages of applying ML techniques, specifically using wrapped feature selection methods such as SHAP values, as a rapid alternative to traditional stepwise covariate modelling^12,14,16^. These methods have shown promise in efficiently identifying relevant covariates without the need for the iterative processes typically associated with stepwise approaches. In this study, we investigated an embedded method for non-linear feature selection, which offers the dual advantage of generating a model and identifying a set of covariates in a single process.

We illustrated the effectiveness of this pipeline using a simple synthetic dataset that included covariates with varying levels of effect: two with low effect (10%) and two with medium effect (30%). Our method successfully identified the relevant covariates, even in the case of high correlation among covariates. Additionally, we introduced a technique for prioritizing covariates by warm-starting the µ-gates and showcased the method’s performance in a scenario involving linear combinations of relevant covariates. We demonstrated that, given sufficient information in the data, relevant covariates will be selected even when penalized. Conversely, favoured covariates will be eliminated if they do not contribute to better data explanation.

Our investigation extended to more complex scenarios to test the robustness of the method. For instance, increasing the IIV to 120%, although the model performed successfully, it made fine-tuning the λ value more difficult. Furthermore, we simulated a population with low-frequency covariates, present in only 5% of the data, which is a challenge for most covariate search method^23^. While the model struggled to identify low-effect covariates in this scenario, it successfully pinpointed medium-effect covariates, demonstrating its potential to detect low-frequency covariates given a sufficient effect size.

To further underscore the advantages of using ML for covariate selection, we simulated an XOR categorical covariate scenario. This exercise illustrated the primary benefit of ML in detecting highly non-linear relationships, which traditional linear-based elimination methods might overlook.

It’s important to note that the conclusions from these challenging scenarios are meant to provide qualitative insights rather than exact quantitative guidance. The exact performance limits, such as the required minimal true effect size or balanced distribution, depend on the overall data quality and the noise within the dataset.

We also compared our embedded method with a combination of XGBoost and SHAP analysis. The results indicated that XGBoost had difficulty in eliminating irrelevant covariates, particularly in the high IIV and highly nonlinear scenarios. In these cases, the SHAP values for non-relevant covariates were similar to those for true covariates, making it challenging to decide which covariates to include. Finally, XGBoost was not able to identify biologically favoured covariates, as it is not possible to set preferred features in XGBoost modelling.

To validate the method outside a controlled simulated environment, we applied it to clinical data from monalizumab and tixagevimab/cilgavimab pooled from several phase III clinical trials.

For monalizumab^21^ the covariates found on clearance and volume of distribution corresponded exactly to the covariates found by the expert. For tixagevimab / cilgavimab^22^, covariates found on three pharmacokinetic parameters: clearance, volume of distribution, and absorption rate constant were a superset of covariates identified by the expert. The primary difference observed was the ‘leakage’ of relevant covariates across all three PK parameters. Thus, we recommend performing a backward selection process after the initial subset of covariates has been identified to mitigate this issue.

The covariates selected by the XGBoost model for the two clinical datasets were largely consistent with those discovered by the SG method.

A significant limitation of our approach is its dependence on Empirical Bayes Estimates, which can become biased and unreliable in conditions of moderate to high shrinkage^24^. Additionally, as demonstrated, in the case of correlation between PK parameters, same covariate can be identified on multiple PK parameters. Consequently, this method should not be viewed as a replacement for the comprehensive model development performed by an expert. Instead, its purpose is to expedite the preselection of covariates in a multivariate context, with the final decision on their inclusion relying on the expertise of pharmacometricians and clinical judgment.

In summary, we demonstrated that a novel machine learning technique developed for covariate selection can significantly accelerate the pre-selection of covariates in popPK model development. It can identify the correct superset of relevant covariates even in challenging scenarios, such as those with low frequency or high correlations. Furthermore, this approach is superior to other recently popularized machine learning methods, such as XGBoost and SHAP, as it can train the model and select covariates in a single process. The identified covariates can be backward eliminated to finalize the model, reducing the time required to build the final model as no forward selection needs to be performed. However, a fundamental limitation of the approach is its reliance on the Empirical Bayes Estimates, which should be addressed in future research.

## Supporting information

Supplementary text with Figures

## Author Contributions

MK and WW performed the analysis; OS and DO prepared the data; MK, AN, SR, CT, DZ and WT wrote the manuscript; MK, OS, AN, RD, DZ, WT and MG designed the research.

