## Supplementary text with Figures for "Stochastic gates for covariate selection in population pharmacokinetics modelling"

### Supplementary material (S1)

#### Details on Stochastic Gates approach

During training, a sample $\epsilon_d \sim N(0, \sigma^2)$ is drawn from a gaussian distribution with fixed standard deviation, and a clipping function $z_d = max(0, \min(1, \mu_d + \epsilon_d))$ is applied before multiplying $z_d$with the input feature $x_d$. Such probabilistic relaxation allows for the joint training of NN parameters and the new parameters, $\mu_d$, using stochastic gradient decent. At the beginning of training $\mu_d$ parameters are initialized around 0, indicating no prior preferences about the feature inclusion.

During inference, $z_d$ is fixed to $z_d = max(0, \min(1, \mu_d))$ to remove stochasticity. The gates have converged if $z_d$is either 0 or 1. In all our experiments we always trained the gates until convergence, however, in case of a weak signal it is possible for the gates to settle at intermediate values.

Loss function, composed of the MSE loss and a penalization of the number of parameters multiplied by coefficient λ) is:

$$\text{Loss} = \frac{1}{N}\sum_{n} \left(Y_{true} - Y_{pred}\right)^2 + \lambda \left|\left|Z\right|\right|_0 \quad . \tag{1}$$

λ is the main hyperparameter governing the number of features used is λ, and needs to be chosen outside of the training procedure. λ that is too large will result in underfitting, while λ that is too small will allow all input features to be used. To ensure the effective learning, both terms in the loss function should be of similar magnitude.

##### *Input data and training procedure*

Before training, input covariates were standardized by dividing them by their standard deviation and shifting them to have a mean value of zero before being processed by the NN. Missing values also must be imputed, and we use standard median/mode values imputation. Categorical features are further encoded using one-hot-encoding: represent each category as a binary vector willed with 0 or 1.
The predicted variable $(\eta_i)$ is also standardized, which is not typically necessary in neural network training. However, this approach facilitates a more consistent selection of the lambda parameter by ensuring that the MSE loss remains on the same scale across different PK parameters.

##### *Training procedure and hyperparameter choice*

Hyperparameter search is performed on the training data only. Train data is divided in 5 non-overlapping folds (so called cross-validation technique), making sure that the studies are equally represented in each fold (patient population and/or dosing strategies can differ between studies). The hyper-parameter tuning will happen in two stages:

1. Coarse search using 4 folds for training and one for validation; where number of layers and neurons, as well as dropout rate and initial lambda parameter will be chosen based on the best validation performance. For this stage, the

configuration involves evaluating various parameters. The number of layers includes options of 1 and 2; the number of nodes per layer comprises 10, 20, 50, or 100 nodes. The Lambda parameter is varied with values set at 0.05, 0.1, 0.5, 1, and 2, while dropout rates of 0 and 0.5 are applied. Optimizer, learning rate, batch size and L2 weight regularization are fixed to Adam, 1e-3, dataset size and 1e-3 respectively.

2. Finer grid-search for lambda parameter using full Cross-Validation strategy. As explained, main parameter to control the number of used features is $\lambda$. To further refine optimal number of features used we perform a grid search using full cross-validation method: for a given lambda parameter we train on 4 folds and calculate performance on the remaining fold. The procedure is repeated 5 times and the mean value of MSE score is calculated, as well as the mean number of used features. Lambda value that gives minimum MSE for minimum (or desired) number of used features is then selected as the optimal one.

In our experiments we performed coarse search only for first (synthetic) dataset and then all hyperparameters except $\lambda$ remained fixed for the rest of the datasets.

On average, across datasets, a complete sequential grid search over the $\lambda$ parameter for a single PK parameter took approximately 30 minutes using an A100 GPU. This process could be further optimized by narrowing the $\lambda$ range—for instance, leveraging Optuna[1] , an automatic hyperparameter optimization algorithm, to identify an optimal $\lambda$ and subsequently testing a smaller range around that value. However, we chose to perform a full grid search to comprehensively illustrate the impact of $\lambda$ on both the loss function and the number of covariates. Alternatively, multiple $\lambda$ values could be evaluated in parallel, as the model requires only ~400MB memory on the GPU.

#### Details on XGBoost and SHAP modelling

To obtain SHAP values for all datasets and targets, we developed XGBoost models for each dataset and target variable. Each dataset was split into training and test sets. Hyperparameter tuning was conducted using RandomizedSearchCV from Python package scikit-learn with a 5-fold cross-validation strategy, exploring a wide range of hyperparameters, including the number of estimators in range [5, 100], learning rate from [0.01, 0.1], maximum depth from [1,3], subsample ratio from [0.2,1], column sample ratio from [0.2,0.6], and regularization parameters such as reg_alpha from [0,1.2], reg_lambda from [10, 40] and min_child_weight from [10, 20] to avoid overfitting. The optimal hyperparameters identified through the random search with 1000 iterations (1000 combinations of the hyperparameters) were then used to train the final XGBoost model. Model evaluation was carried out by predicting on both training and testing sets, with performance metrics such as MSE and $R^2$ score being recorded. SHAP values were utilized to interpret the model's predictions and visualize feature importance, with SHAP summary plots being generated for each dataset and target variable. Results are shown below.

Since XGBoost does not require standardization of the output variable, comparing the MSE loss between rescaled and non-rescaled variables is not feasible. Instead, we used the $R^2$ metric for comparison, as it is invariant to rescaling of the output variable.

#### XOR example for nonlinear function

To exploit the power of machine learning versus more traditional GAM models we simulate a covariate interaction using a well-known nonlinear XOR-like structure:

$$CL_k = CL_{pop} \times \left(1 + \theta \times (\mathrm{CAT}_{\mathrm{ik}} - CAT_{jk})^2\right) \qquad (1)$$

As a proof of concept, we use SMK and COPD as binary covariates, although this setup is not intended to reflect a realistic biological relationship. In this construction, the individual effect of each covariate appears weak or nonexistent when examined independently. Traditional univariate analyses or models such as linear regression and GAMs fail to capture the dependency of the PK parameter ($CL_k$) on either SMK or COPD alone.

The true interaction only becomes apparent when SMK and COPD are considered jointly. The relationship follows an XOR-like pattern: $CL_k$ varies meaningfully only when the two covariates differ (i.e., one is 0 and the other is 1). This creates a nonlinear, non-additive structure that cannot be adequately modeled by linear terms or additive smooth functions, underscoring the need for models that can capture complex interactions—such as machine learning methods.

##### *Covariates abbreviation:*

- BHT (baseline height),
- BWT (baseline weight),
- BBILI (baseline bilirubin),
- BSCR (baseline serum creatinine),
- BBSA (baseline body surface area),
- BALB (baseline albumin),
- BALT (baseline alanine aminotransferase),
- BAST (baseline aspartate aminotransferase),
- BCRCL (baseline creatine clearance),
- BEGFR (baseline estimated glomerular filtration rate),
- CLD (chronic liver disease),

- SMK (smoking status),
- COPD (chronic obstructive pulmonary disease)
- CKD (chronic kidney disease)

#### Figures:

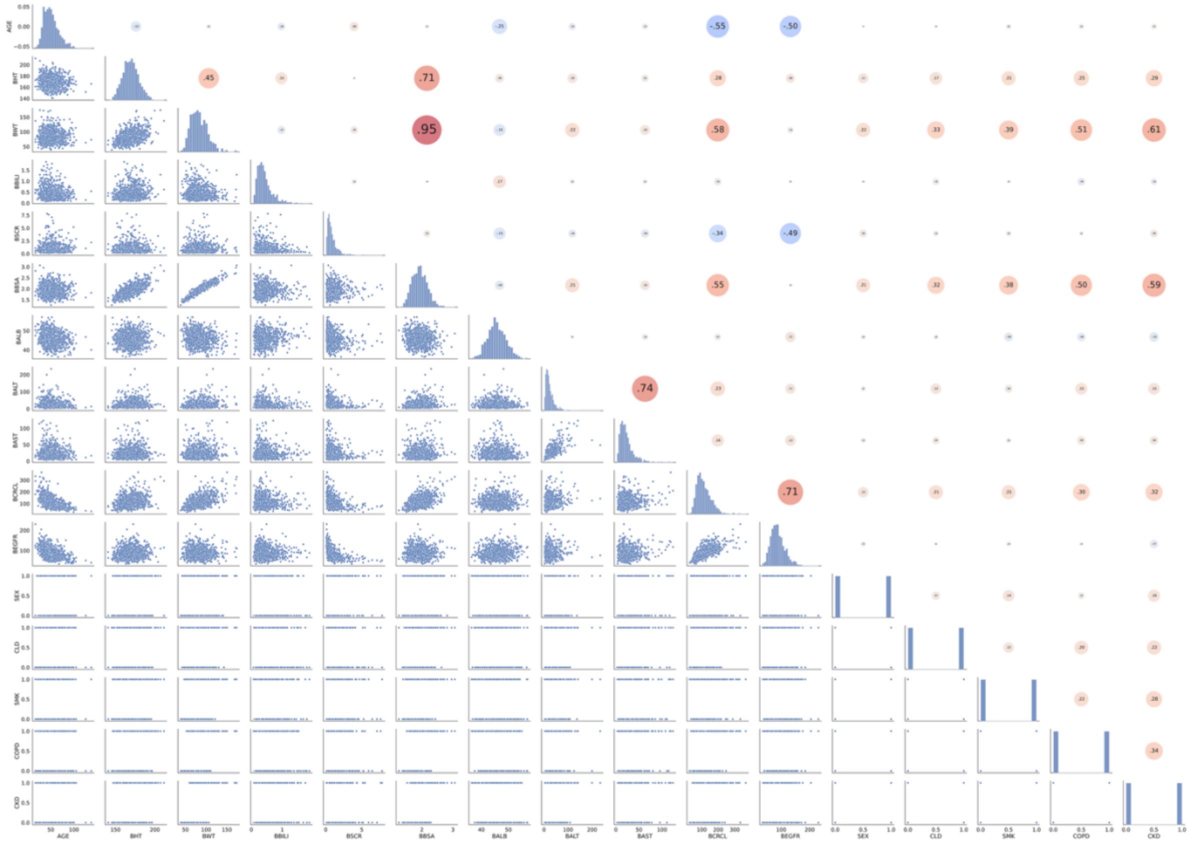

*Figure S1: Synthetic dataset covariates correlations. Covariates are AGE, BHT (baseline height), BWT (baseline weight), BBILI (baseline bilirubin), BSCR (baseline serum creatinine), BBSA (baseline body surface area), BALB (baseline albumin), BALT (baseline alanine aminotransferase), BAST (baseline aspartate aminotransferase), BCRCL (baseline creatine clearance), BEGFR(baseline estimated glomerular filtration rate), SEX, CLD (chronic liver disease), SMK (smoking status), COPD (chronic obstructive pulmonary disease) and CKD (chronic kidney disease)*

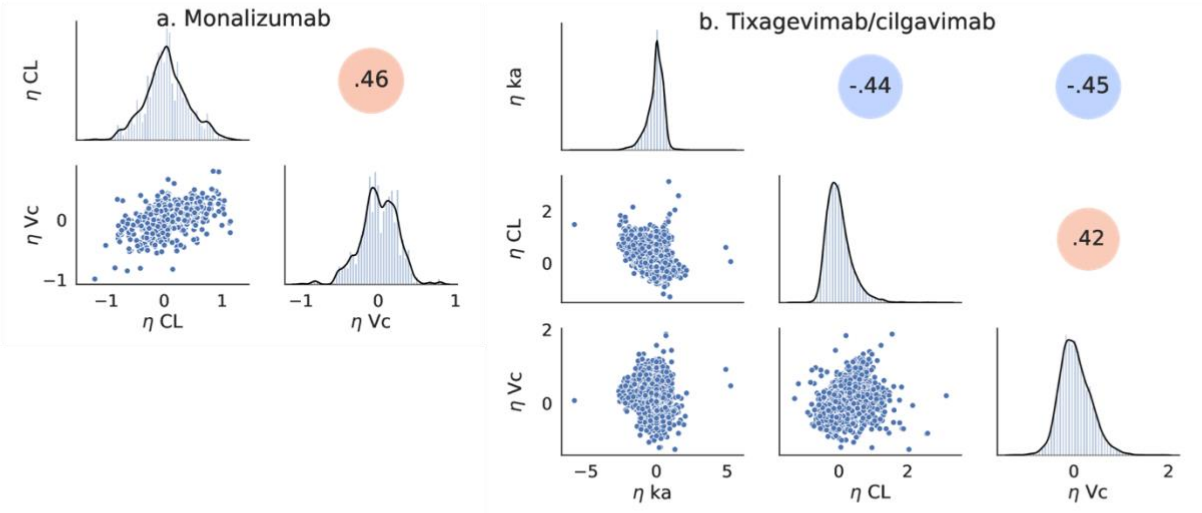


*Figure S2: ETA distribution for monalizumab (a) and tixagevimab / cilgavimab (b)*

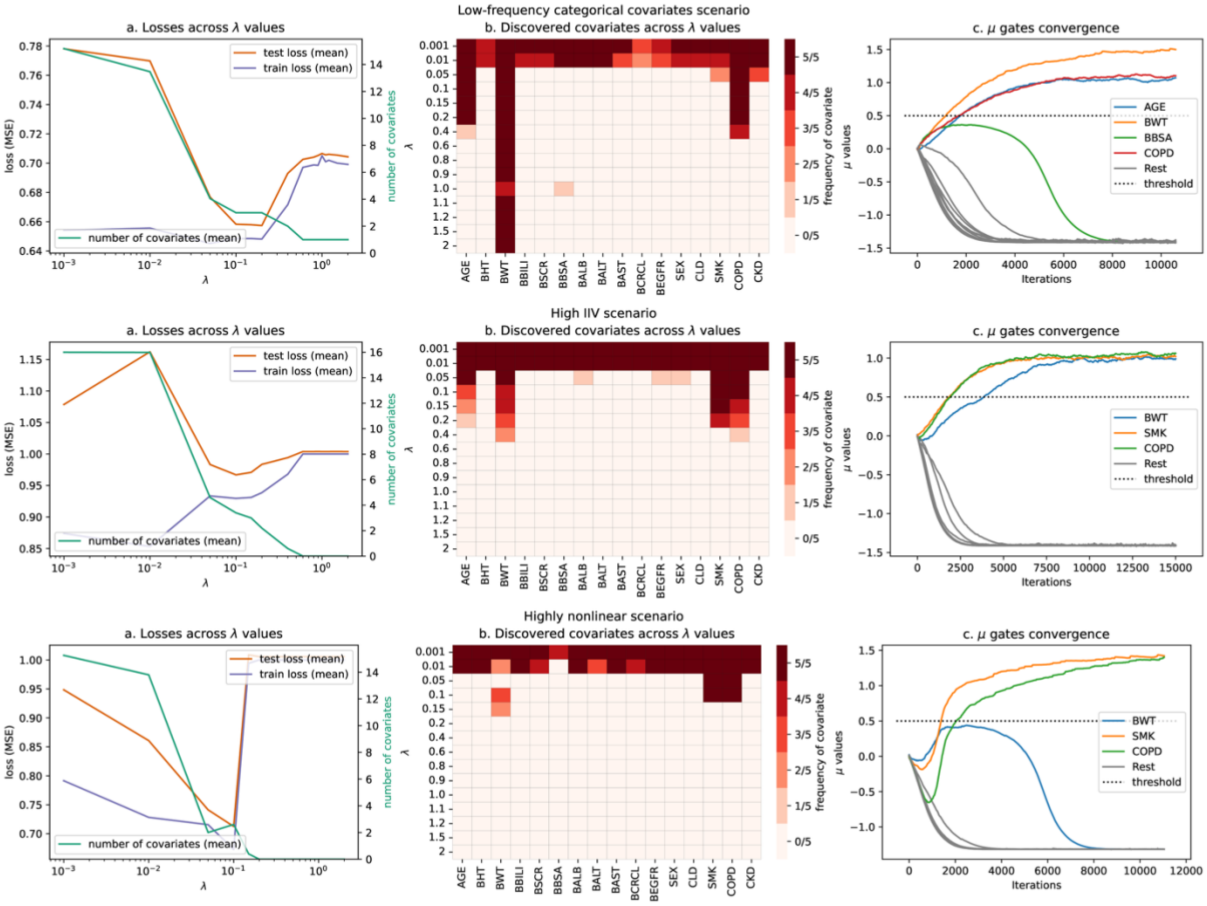


*Figure S3: Stochastic gates output for low frequency scenario (top), high IIV scenario (middle), and highly non-linear XOR scenario (bottom).*

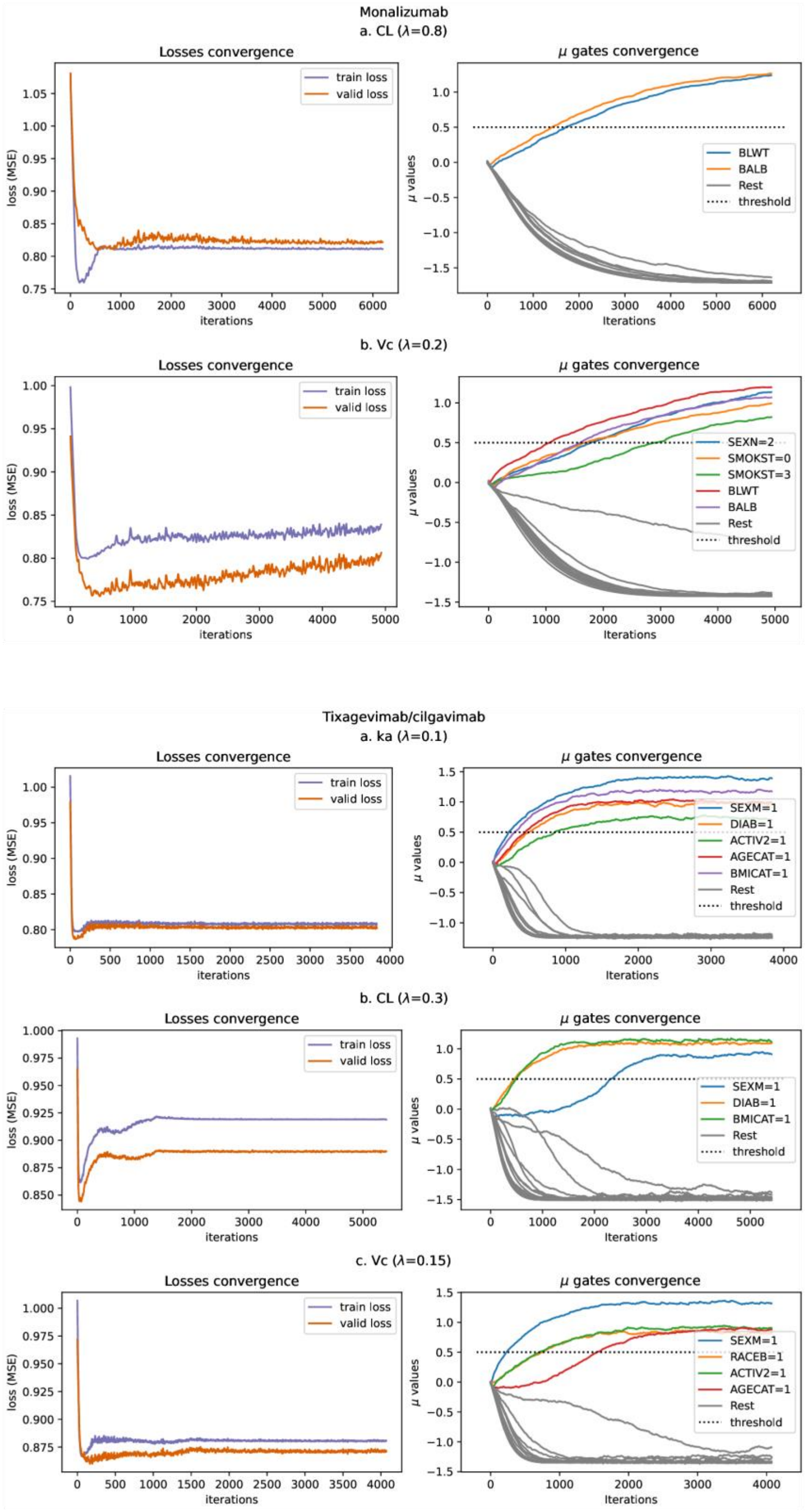


*Figure S4: Final train/test split outputs. Top: monalizumab — (a) CL and (b) Vc parameters. Bottom: tixagevimab/cilgavimab — (a) ka, (b) CL, and (c) Vc parameters. Left panels show loss convergence; right panels show μ-gates convergence.*

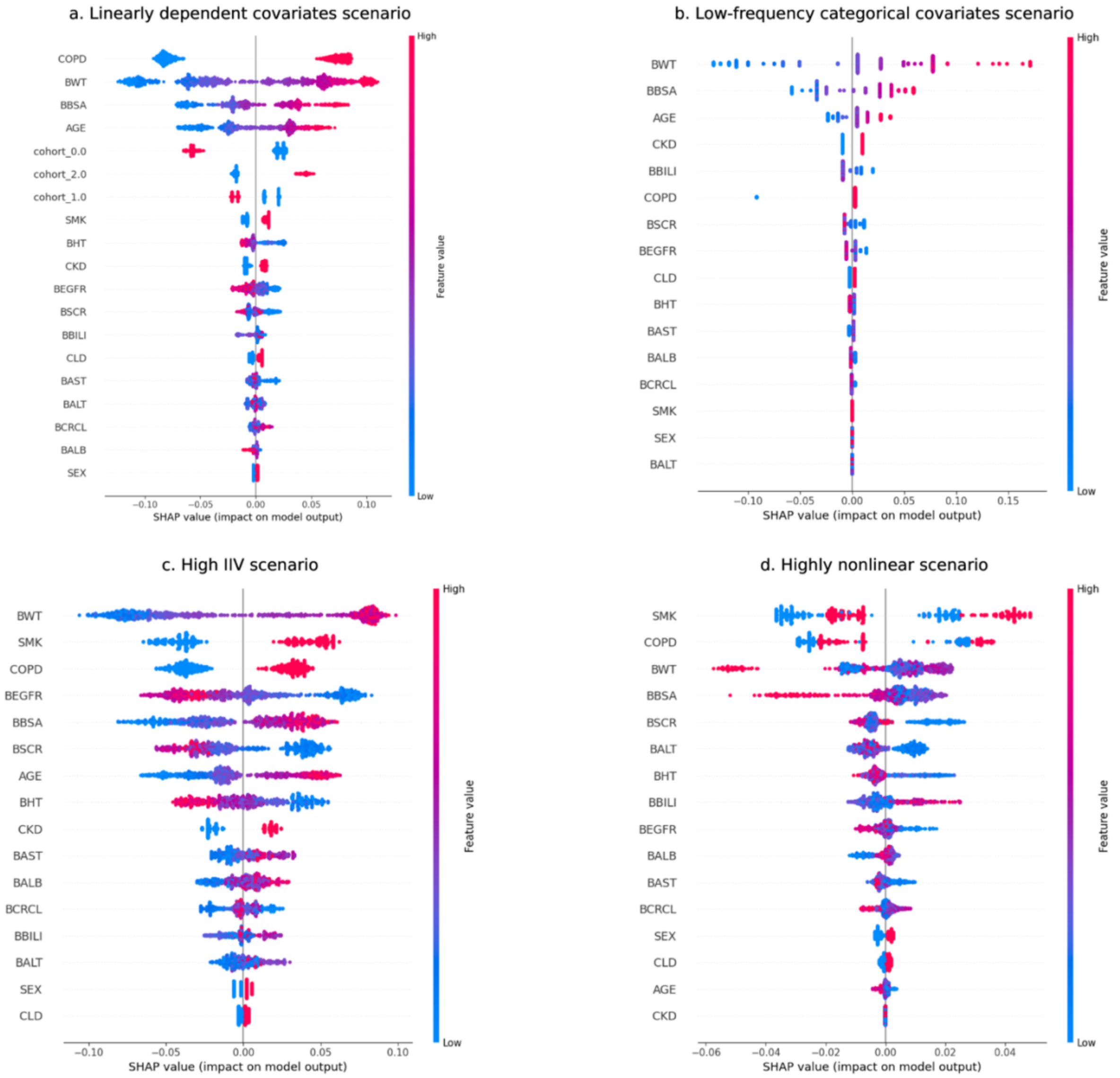


*Figure S5: XGBoost-obtained SHAP values for (a) Linearly dependent, (b) Low-frequency, (c) High IIV, (d) Highly nonlinear scenarios*

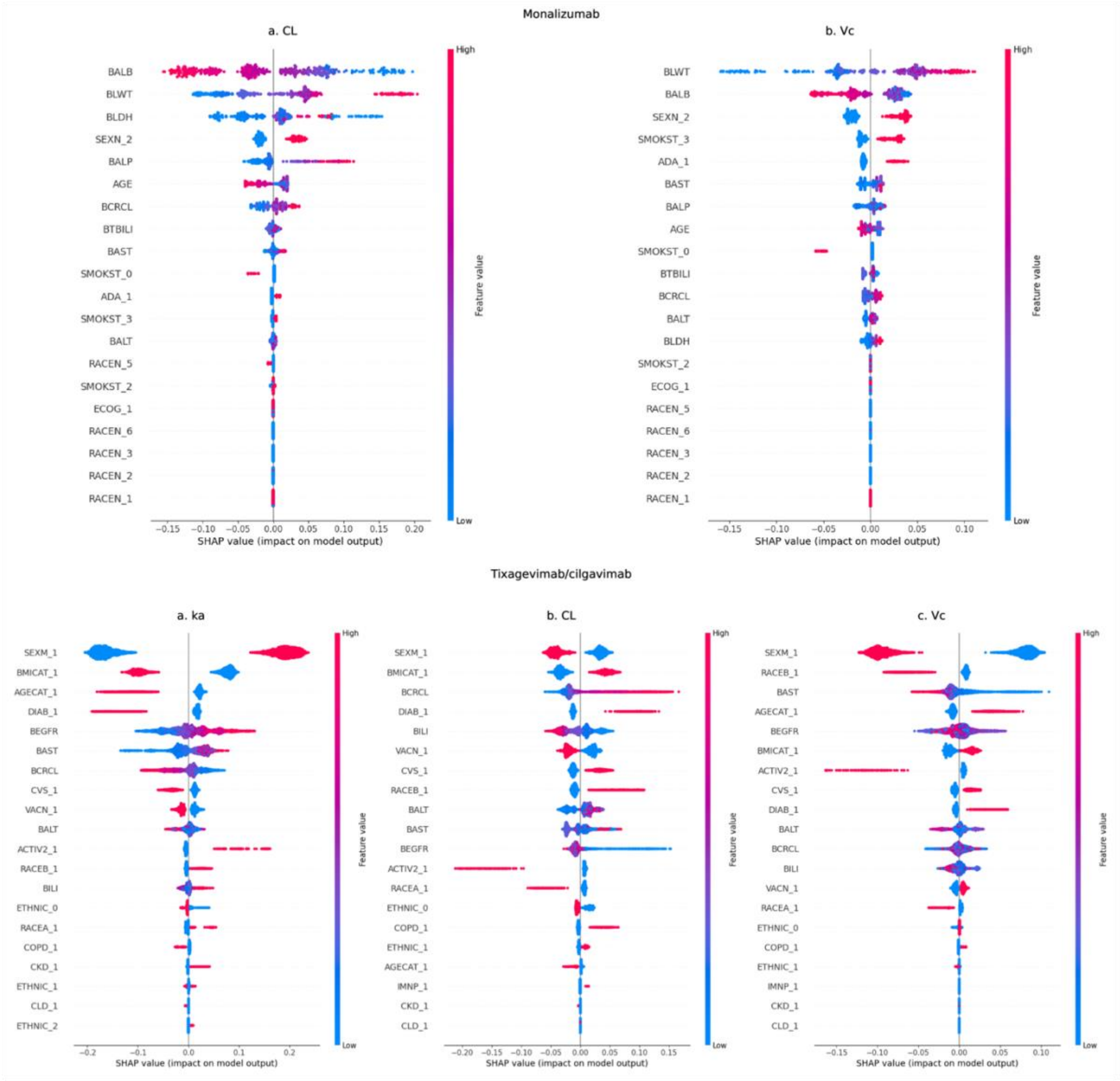


*Figure S4: SHAP values from the XGBoost model. Top: monalizumab — (a) CL and (b) Vc parameters. Bottom: tixagevimab/cilgavimab — (a) ka, (b) CL, and (c) Vc parameters.*

#### Tables

*Table S1 Performance comparison of NN with SG and XGBoost regression on all datasets*

| **Dataset name** | **Train/Test R2 with NN with SG** | **Train/Test R2 with XGBoost** |
|---|---|---|
| Reference | 0.58/0.61 | 0.63/0.61 |
| Linearly dependent covariates | 0.58/0.61 | 0.63/0.61 |
| Low-frequency categorical covariates | 0.35/0.4 | 0.38/0.36 |
| High IIV | 0.06/0.1 | 0.09/0.07 |
| Highly nonlinear | 0.32/0.2 | 0.32/0.21 |
| monalizumab CL | 0.19/0.16 | 0.37/0.11 |
| monalizumab Vc | 0.17/0.19 | 0.31/0.11 |
| tixagevimab / cilgavimab CL | 0.08/0.08 | 0.20/0.16 |
| tixagevimab / cilgavimab Vc | 0.12/0.10 | 0.19/0.11 |
| tixagevimab / cilgavimab ka | 0.19/0.18 | 0.25/0.20 |
